# Heat stress disrupts early development and photosymbiosis in *Cassiopea* jellyfish

**DOI:** 10.1101/2025.04.19.649683

**Authors:** Celeste Robinson, Jingchun Li, Ruiqi Li, Viridiana Avila-Magaña

## Abstract

Photosymbioses between Cnidarians and algae are widespread in marine ecosystems. The jellyfish *Cassiopea*-*Symbiodinium* symbiosis serves as a valuable model for studying host-symbiont interactions in photosymbiotic organisms. Despite its ecological similarity to coral symbiosis, the effects of rising sea surface temperatures on *Cassiopea* symbiosis, particularly during early developmental stages, remain unexplored. By exposing *Symbiodinium* cultures to heat stress and subsequently using these symbionts to colonize jellyfish polyps under ambient and elevated temperature conditions, we study the impact of heat on microbe-stimulating metamorphosis. We observed a significant reduction in chlorophyll concentration in heat-stressed *Symbiodinium* algae. Polyps colonized with these symbionts exhibited delayed metamorphosis under ambient conditions and failed to undergo metamorphosis under continued heat stress. Additionally, we found abnormal ephyra morphology and increased rates of asexual reproduction under heat stress. Our findings suggest that ocean warming may disrupt critical stages of *Cassiopea* metamorphosis and development by impairing symbiosis, ultimately threatening their population stability under warming marine environments.

## Introduction

Photosymbiotic relationships between marine invertebrates and photosynthetic algae play a crucial role in tropical marine ecosystems, driving primary production, nutrient cycling, and biodiversity (1). Among these, cnidarian photosymbiosis is particularly significant, as it forms the foundation of one of the most productive habitats on Earth—coral reefs. However, rising sea surface temperatures due to climate change threaten these associations by disrupting symbiotic interactions (2). Understanding the effects of thermal stress on these partnerships is essential for predicting broader ecological consequences. In recent years, research has largely focused on corals due to their vulnerability to climate change, but the impacts of heat stress on other photosymbiotic cnidarians remain understudied.

Among other photosymbiotic cnidarians, the upside-down jellyfish, *Cassiopea xamachana*, is a particularly interesting model due to its symbiosis with Symbiodiniaceae dinoflagellates, which not only provide energy but also regulate its metamorphosis. *C. xamachana* has a complex life cycle involving both sexual and asexual reproduction (3). Polyps reproduce asexually through stolon budding, generating planuloid buds (4), while sexually mature medusae produce planula larvae that settle and develop into polyps (5). Unlike most jellyfish, where sexual reproduction is influenced by seasonal temperature changes (6), *C. xamachana* requires colonization by Symbiodiniaceae symbionts to initiate metamorphosis. Without these symbionts, polyps remain indefinitely at the benthic stage and fail to undergo strobilation (5). Once infected, polyps begin strobilation, producing ephyrae that develop into adult medusae (Fig 1).

**Fig 1.**
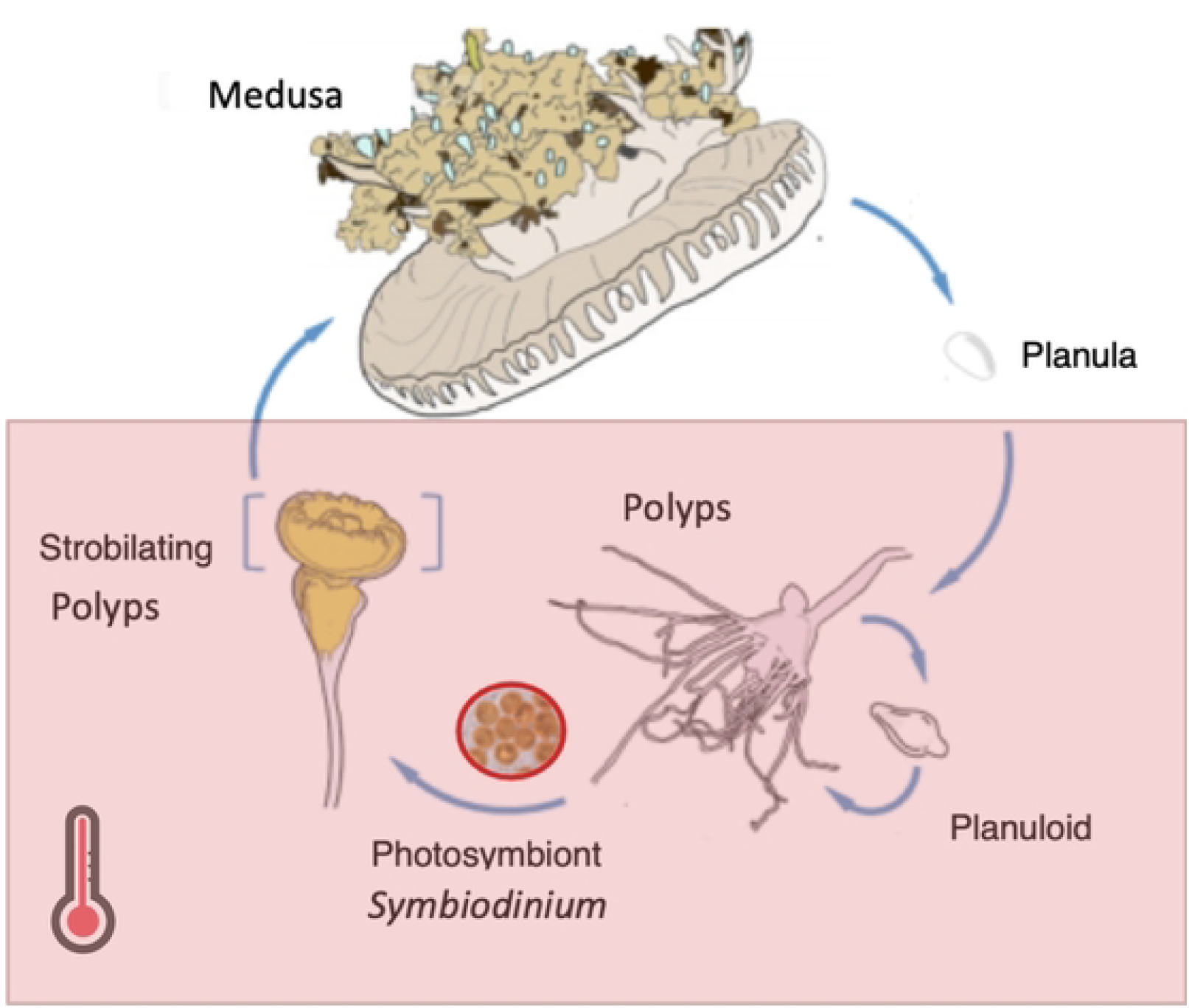
life cycle of *Cassiopea xamachana*. *C. xamachana* can reproduce sexually and asexually, with strobilation induced by Symbiodinium Colonization. Polyps can regenerate after strobilation. The red box highlights the life stage examined in this study. Figure was modified from Ohdera 2018.

Relying on symbionts for metamorphosis may make *C. xamachana* particularly vulnerable to rising temperatures. Although its primary symbiont, *Symbiodinium microadriaticum*, is relatively thermotolerant compared to other Symbiodiniaceae species (7), prolonged heat stress can still lead to pigment loss, photosynthetic dysfunction, and even photoinhibition in *S. microadriaticum* (8,9). In corals, *Acropora digitifera* larvae exhibited higher mortality or prematurely underwent metamorphosis under heat stress (10). The impact of heat stress on *C. xamachana* development remains unexamined, but studies have shown that ocean acidification can induce developmental abnormalities, such as causing an inverted bell shape (11). Investigating how heat stress affects *C. xamachana*’s reproduction and development is particularly valuable, as it provides insights into both the consequences of nutritional symbiosis breakdown and the cascading effects of heat stress on metamorphosis cues.

In this study, a heat-stress experiment was conducted on *C. xamachana* polyps and their symbiont, S. *microadriaticum*, to examine how elevated temperatures influence their symbiotic relationship, particularly in terms of development and reproduction. A combination of morphological observations, symbiont density quantification, and chlorophyll concentration analysis was used to quantitatively and qualitatively assess the effects of heat stress on *C. xamachana* metamorphosis and reproduction. By addressing these questions, this study provides insights into how ocean warming may disrupt cnidarian-algal symbiosis and impact *C. xamachana* populations in a warmer ocean.

## Materials and methods

### Materials

*C. xamachana* polyps were originally collected from Key Largo, Florida Keys. A clonal polyp line was established in Mónica Medina’s lab at Pennsylvania State University. A subculture was later established at the University of Colorado Boulder and used in this study. These clonal polyps were maintained in filtered artificial seawater tanks at a constant temperature of 26°C, and a salinity of 35 ppt. Polyps were fed every other day with the San Francisco strain of *Artemia* brine shrimp. The KB8 strain of *Symbiodinium microadriaticum* (12) was cultured in 0.1 M F/2 media. Cultures were maintained in a temperature and light-controlled incubator (Percival Scientific, Perry, USA) at 26°C, 140 μmol quanta m−^2^s−^1^, under a 12:12 h light-dark cycle, at a density of ~4.26 million cells/mL.

### Experiment design

To assess the impact of thermal stress on *C. xamachana* development and reproduction, we established three experimental groups using *S. microadriaticum* to infect *C. xamachana* polyps (Fig 2). A temperature of 34°C was chosen for heat treatment because that is the upper limit of relevant ecological broad temperature range (26–34°C) for *C. xamachana* (13,14), and it has been reported to support the maintenance of homeostatic bell pulsation within that range (15). Symbionts were pre-exposed to 34°C for 8 days before colonization, and chlorophyll content was measured on days 6 and 8 using a NanoDrop spectrophotometer (Thermo Scientific, Waltham, USA) following Klein et al. (2019) (16). In addition, algal cell density was quantified using a Countess II FL automated cell counter (Invitrogen, Waltham, USA). Colonizations were conducted with symbionts in their exponential growth phase at 1 × 10^6^ cells ml^−1^ following Medina et al. 2021(17). The heat group was infected with symbionts cultured at 34°C and maintained at 34°C. In the recovery group, polyps were infected with symbionts cultured at 34°C but maintained at 26°C after colonization. The control group consisted of polyps and symbionts both raised and maintained at 26°C. Colonizations were conducted in three 6-well plates, one per experimental group, with about 50 polyps per plate for replication (Fig 2). 12 hours post colonization, we removed all water from the 6 well plates and rinsed them, replenishing the wells with filtered sea water and resumed normal feeding and cleaning procedures.

**Fig 2.**
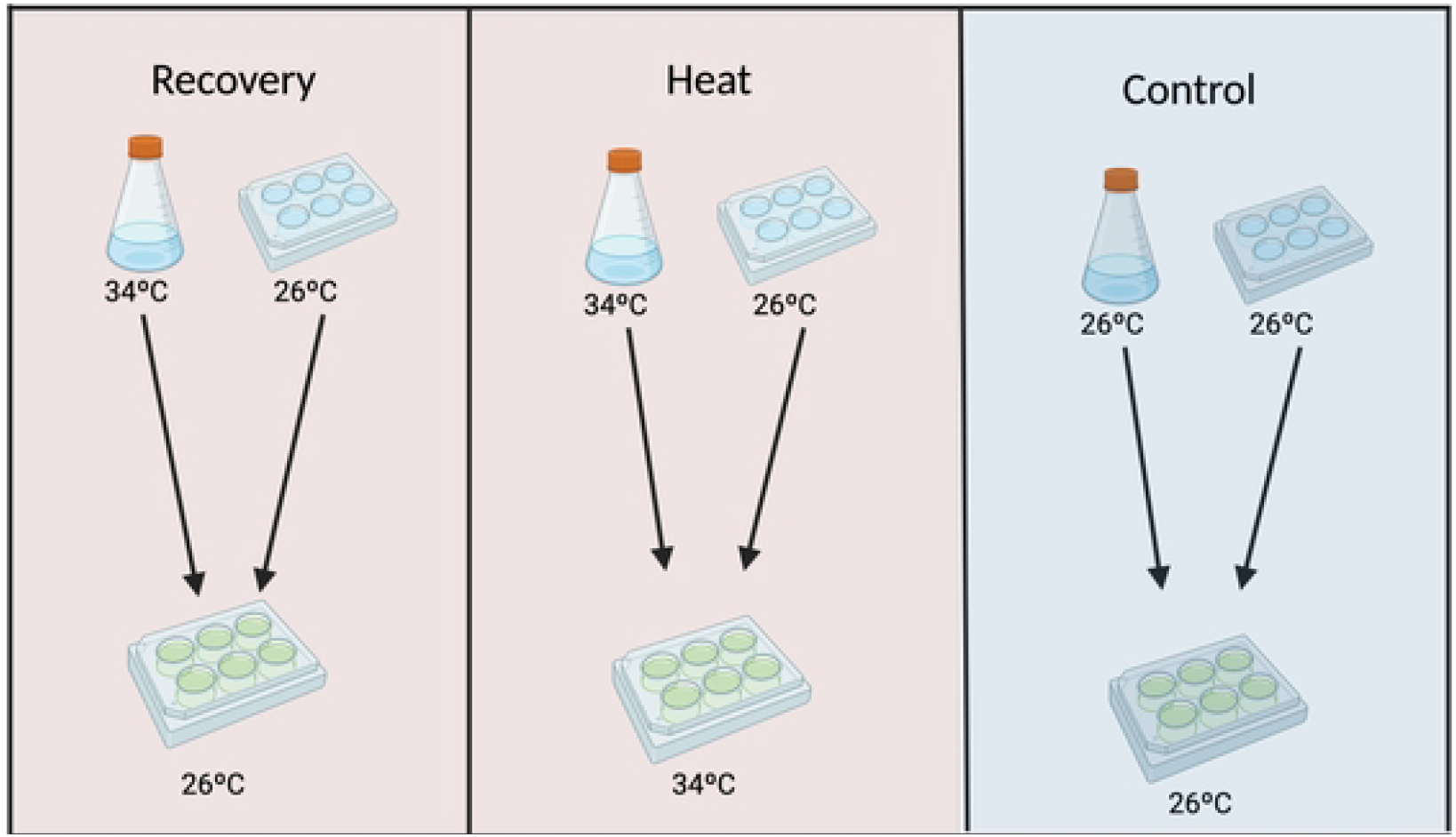
Experimental design to assess the impact of thermal stress on *C. xamachana* development. Three groups were used: 1) Heat – polyps infected with symbionts cultured at 34°C and maintained at 34°C; 2) Recovery – polyps infected with symbionts cultured at 34°C but maintained at 26°C; and 3) Control – polyps and symbionts both cultured and maintained at 26°C.

### Measurements

To assess the impact of heat stress on host asexual reproduction, the number of planuloid buds in each well was recorded throughout the experiment. Photos of ephyrae were captured using a Micromaster M002 microscope (Fisher Scientific, Waltham, USA) to identify potential morphological abnormalities. The number of polyps undergoing strobilation was counted every 3– 5 days to evaluate the effect of heat stress on metamorphosis.

To quantify algal cell density within the ephyrae, five ephyrae from the control group and ten from the recovery group, including five inverted and five normal ephyrae, were collected fifteen days after the onset of strobilation. Samples were flash-frozen, then thawed, macerated with a pestle until homogenized, and vortexed for 20 minutes. The homogenized suspension was loaded onto a hemocytometer, and algal cells were counted under a Fisher Scientific Micromaster M002 microscope at 10× magnification. At least four replicates were counted per sample.

### Statistical analyses

The effect of heat stress and morphological forms on *in hospite* algal cell density was analyzed by comparing algal cell density in ephyrae raised under normal temperatures to those exposed to heat stress (normal and inverted forms). T-tests were performed to evaluate statistical significance between groups using R (18). Plots were generated using ggplot2 package in R.

## Results and discussion

### Heat-stressed *Symbiodinium* prevent *Cassiopea* strobilation

Strobilation cues in jellyfish vary widely among species and can be influenced by environmental factors such as temperature, salinity, and light exposure. For example, *Aurelia aurita* strobilates after prolonged cold periods (19), whereas species like *Cephea cephea* and *Rhopilema verrilli* respond to elevated temperatures (20,21). Unlike these species, *Cassiopea* relies primarily on colonization by endosymbiotic *Symbiodinium* to initiate strobilation (5). There is some evidence suggesting that metabolic products derived from symbiont, specifically carotenoids may be important to initiate *C. xamachana* development (13), although it is unclear whether there are additional metabolites or a combination of other factors regulating and triggering host development.

Our experiments revealed a significant delay in symbiont-induced strobilation when symbionts were pre-exposed to thermal stress in the recovery group. While control polyps completed strobilation by day 19, full strobilation in the recovery group was delayed until day 36 (Fig 3). In contrast, no strobilation occurred under continuous heat stress. Although it is possible that *Cassiopea* is inherently unable to strobilate under prolonged heat stress, the significant delay observed in the recovery group suggests that physiological changes in the symbionts are more likely to be responsible for the disruption in metamorphosis. While symbiont cell viability did not show a significant difference (Fig 4a), chlorophyll a quantification indicated a substantial reduction in pigment concentration under thermal stress (Fig 4b), with symbionts appearing visibly lighter in color compared to the darker brown control cultures (Fig 4c). These findings suggest that heat stress alters symbiont physiology in a way that impairs their ability to induce timely strobilation in *Cassiopea*.

**Fig 3.**
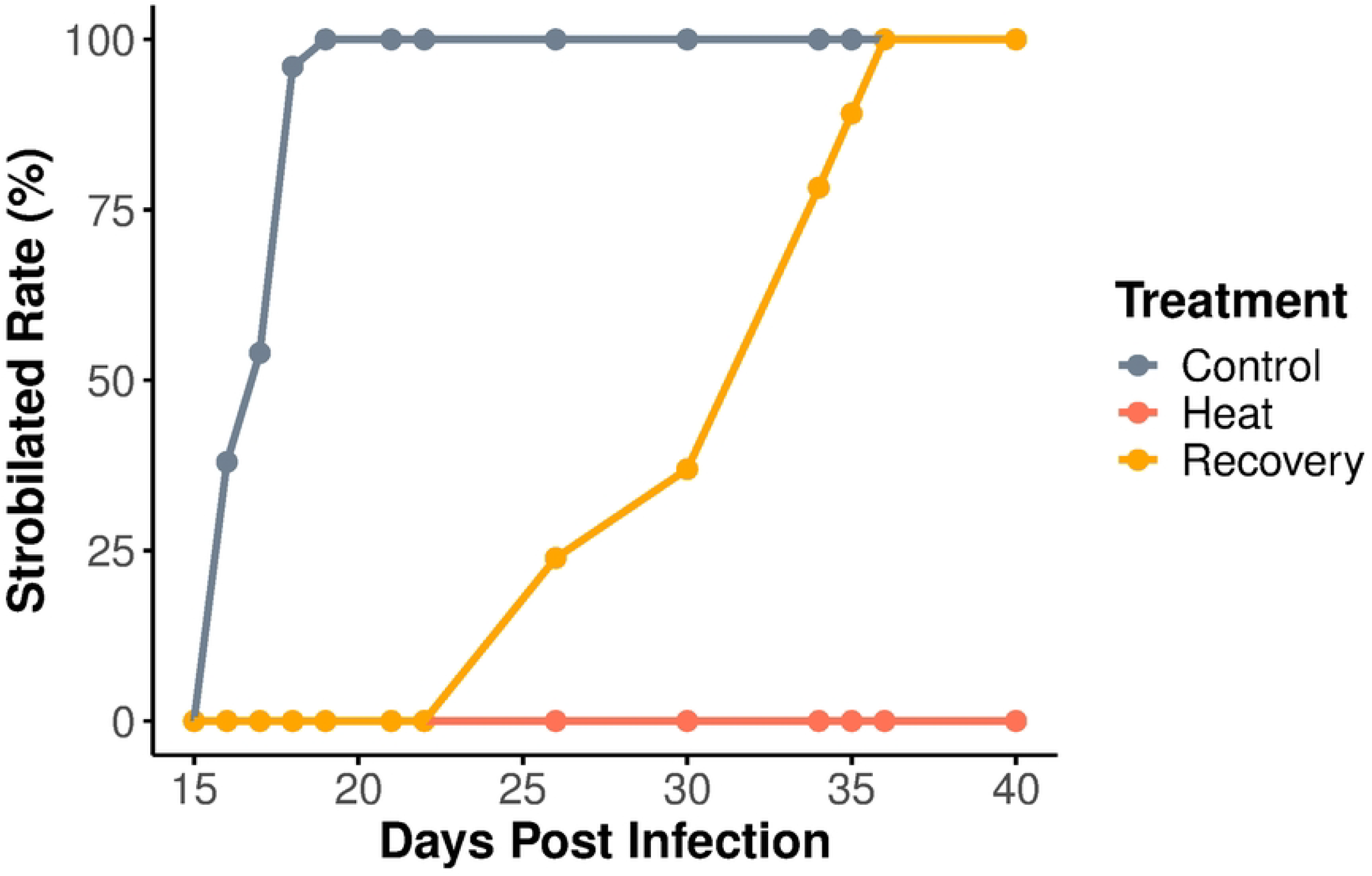
Percentage of polyps undergoing strobilation over time in each treatment group. Control group completed strobilation by day 19, while strobilation in the recovery group was delayed until day 36. No strobilation was observed in the heat group. N=50.

**Fig 4.**
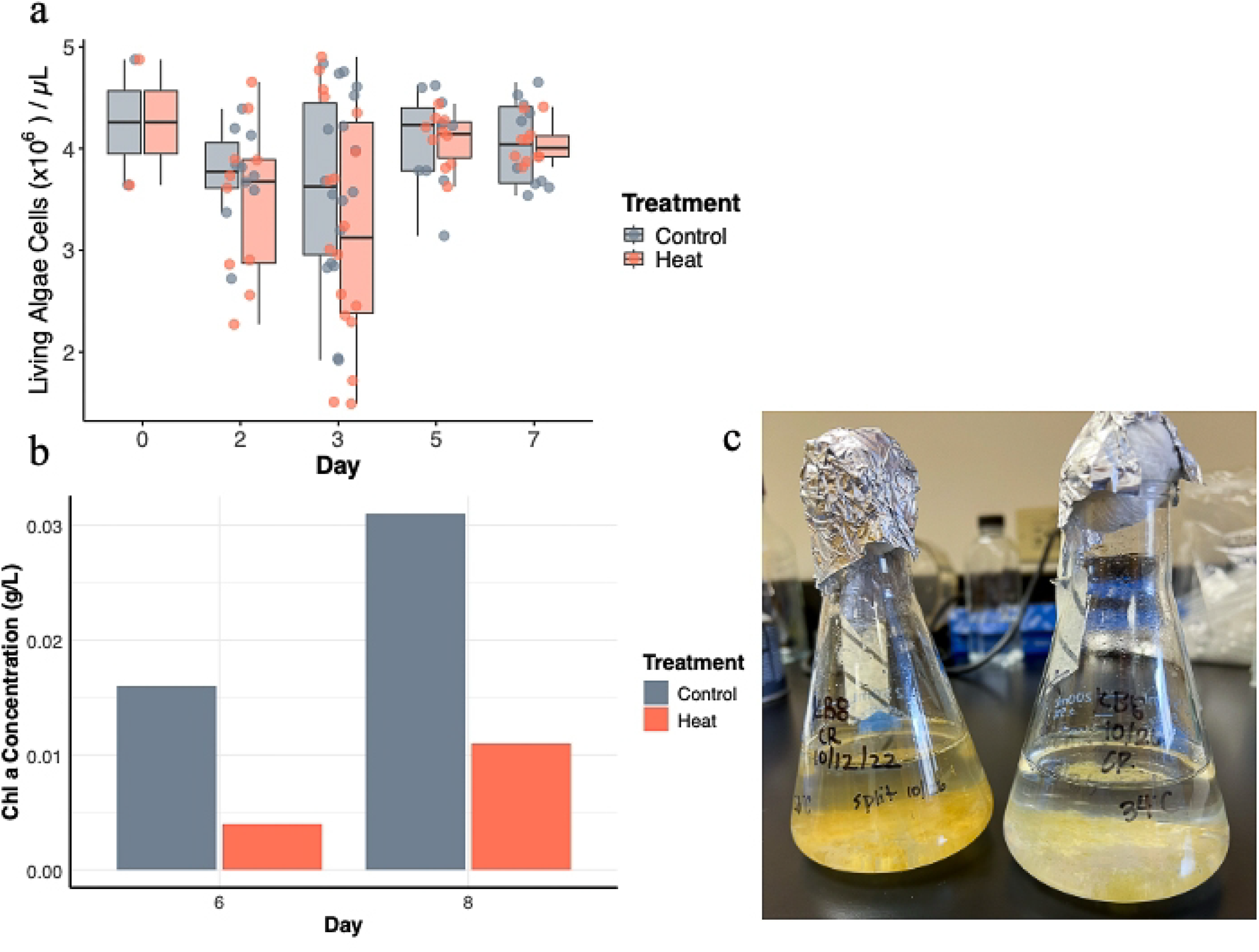
Effects of heat stress on *S. microadriaticum*. (a) Symbiont cell density did not differ significantly across treatments. (b) Chlorophyll content was significantly reduced under thermal stress. (c) Symbionts exposed to heat (right) appeared visibly lighter in color compared to the darker brown control cultures (left).

The impact of thermal stress on *Symbiodinium* is complex, affecting various physiological and molecular processes. Transcriptomic studies have revealed widespread but subtle gene expression changes in *Symbiodinium* under thermal stress, affecting antioxidant networks, chaperone proteins, photosynthesis activities, and metabolic pathways (22). Heat stress can also lead to photoinhibition by impairing photosystem II (PSII) (9). However, impaired photosynthesis alone is unlikely to be the direct cause of delayed or halted strobilation in *Cassiopea*. Since carotenoid metabolites are believed to play a role in inducing *Cassiopea* strobilation (13) and previous studies have shown that thermal stress leads to a loss of pigments in the Light Harvesting Complex, resulting in reduced chlorophyll A and carotenoid levels in *Symbiodiniaceae* (23), the observed loss of symbiont pigmentation may explain the effect of heat stress on strobilation. Other studies have shown that indoles can induce strobilation in aposymbiotic *Cassiopea andromeda* polyps, suggesting that certain chemical compounds may serve as direct triggers for *Cassiopea* strobilation (19,21). Metabolomic studies on *Symbiodiniaceae* under heat stress have identified distinct metabolic changes (25). However, further research on the *Cassiopea* holobiont is necessary to pinpoint additional photosymbiont metabolites that contribute to strobilation.

### Alternated algal density and developmental abnormalities in ephyrae with symbionts exposed to heat

While photosynthesis may not be the direct cue for host metamorphosis, it is essential for *Cassiopea* development. Aposymbiotic *Cassiopea* polyps induced to strobilate by chemical cues often exhibit developmental abnormalities, including reduced ephyra size, malformed sensory organs, crimped marginal lappets, delayed polyp recovery after strobilation, or, in some cases, death (21). Our results align with these findings, as 90% (45 out of 50) of ephyrae from the recovery group exhibited an inverted bell shape and were significantly smaller (~0.3 mm) compared to control ephyrae (~0.5–0.6 mm) (Fig 5a).

**Fig 5.**
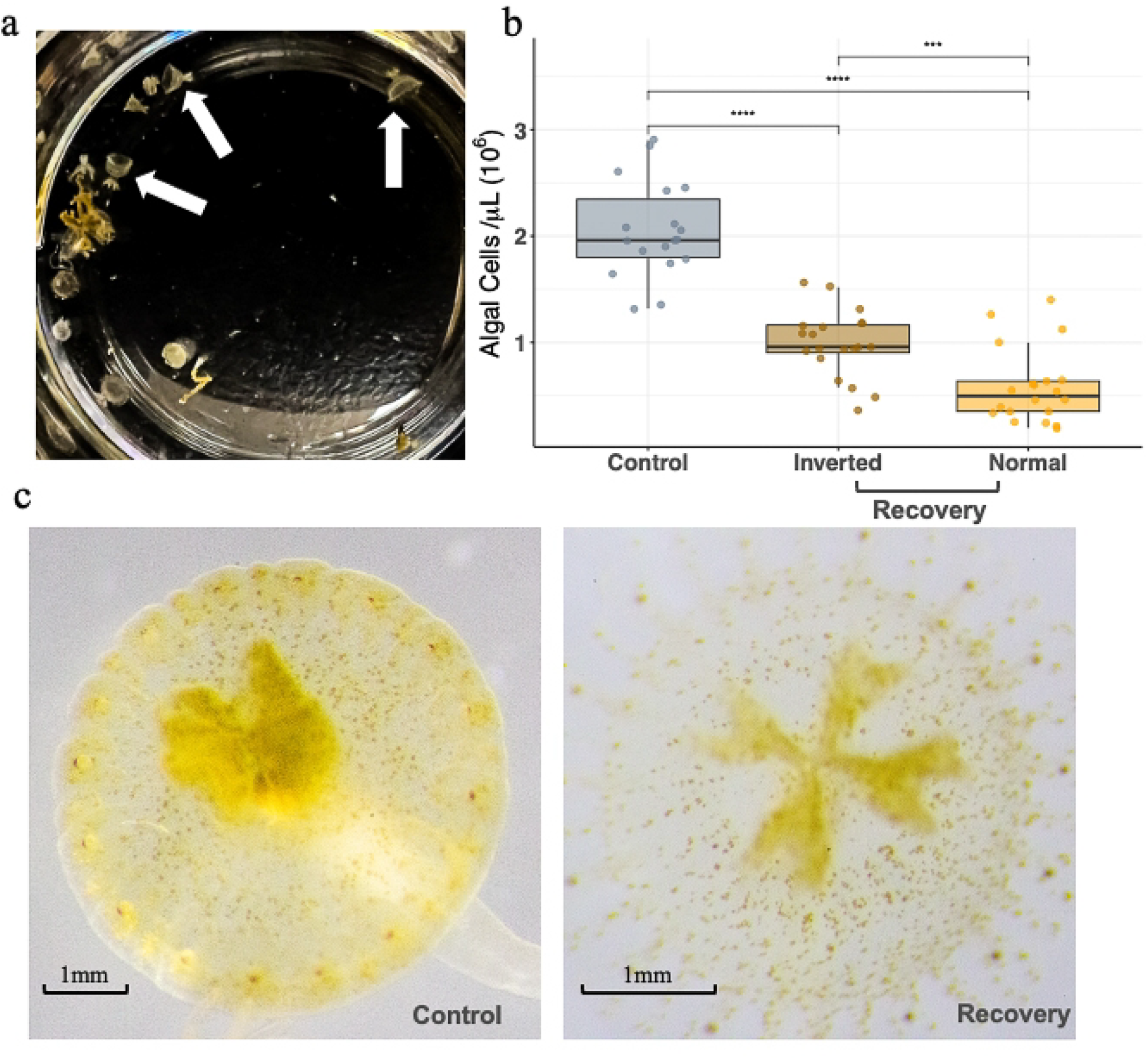
Effects of heat stress *on C. xamachana* developmental morphology and symbiont density. a) Most ephyrae in the recovery group displayed an inverted bell shape as shown in the photo. b) Inverted ephyrae had significantly higher symbiont densities than normal-shaped ephyrae within the recovery group, notice that both were lower than in the control group. c) Visual comparison shows reduced symbiont density in ephyrae from the recovery group compared to the control.

The exact cause of *Cassiopea* heat-stressed ephyra developmental abnormalities remains unknown. However, ephyrae with an inverted bell shape contained significantly more symbiont cells than normal-shaped ephyrae within the recovery group (p < 0.01, Fig 5b). Both inverted and normal ephyrae in the recovery group had significantly lower algal densities compared to those in the control group, this was further confirmed by visual inspection (p < 0.001, Fig 5bc).

Why do inverted ephyrae have a higher symbiont density than normal-shaped ones in the recovery group? Heat stress can alter the host-symbiont relationship, potentially leading to a breakdown in mutualism. Under certain conditions, symbionts proliferate more rapidly within the host and exhibit higher dispersal rates while simultaneously reducing host growth and reproduction (26).

Thermal stress has been documented as a disruptor of the host development and immune system (27), which ultimately leads to uncontrolled symbiont proliferation and perhaps to a shift from a mutualistic to parasitic relationship. Further investigation is needed to support this, since we did not directly measure host fitness. Similar inverted bell-shaped abnormalities have also been observed in *Cassiopea* raised under severe seawater acidification (pH = 7.0) (11). Further research is needed to investigate the underlying mechanisms driving these developmental abnormalities and their broader ecological implications.

### Increased asexual reproduction in polyps infected with heat-Stressed algae

Continuous heat stress prevents *Cassiopea* polyps from strobilating, but does not alter their ability to reproduce asexually. During the polyp stage, *Cassiopea* can still reproduce asexually through budding (29). In our experiment, polyps infected with heat-stressed algae exhibited an increase in planuloid bud production compared to the control group, with both heat and recovery groups showing a higher overall number of buds (Fig 6). Jellyfish reproduction often follows a boom-and-bust cycle (30), where rapid increases in reproductive output are followed by population crashes. The underlying mechanisms remain unclear, but it is possible that *C. xamachana* polyps under heat stress allocate their limited energy and nutrients toward offspring production as a survival strategy. Similar stress-induced reproductive responses have been observed across the animal kingdom when resources are scarce or environmental pressures intensify (31), suggesting that increased planuloid budding may be a stress response triggered by physiological changes from thermal stress.

**Fig 6.**
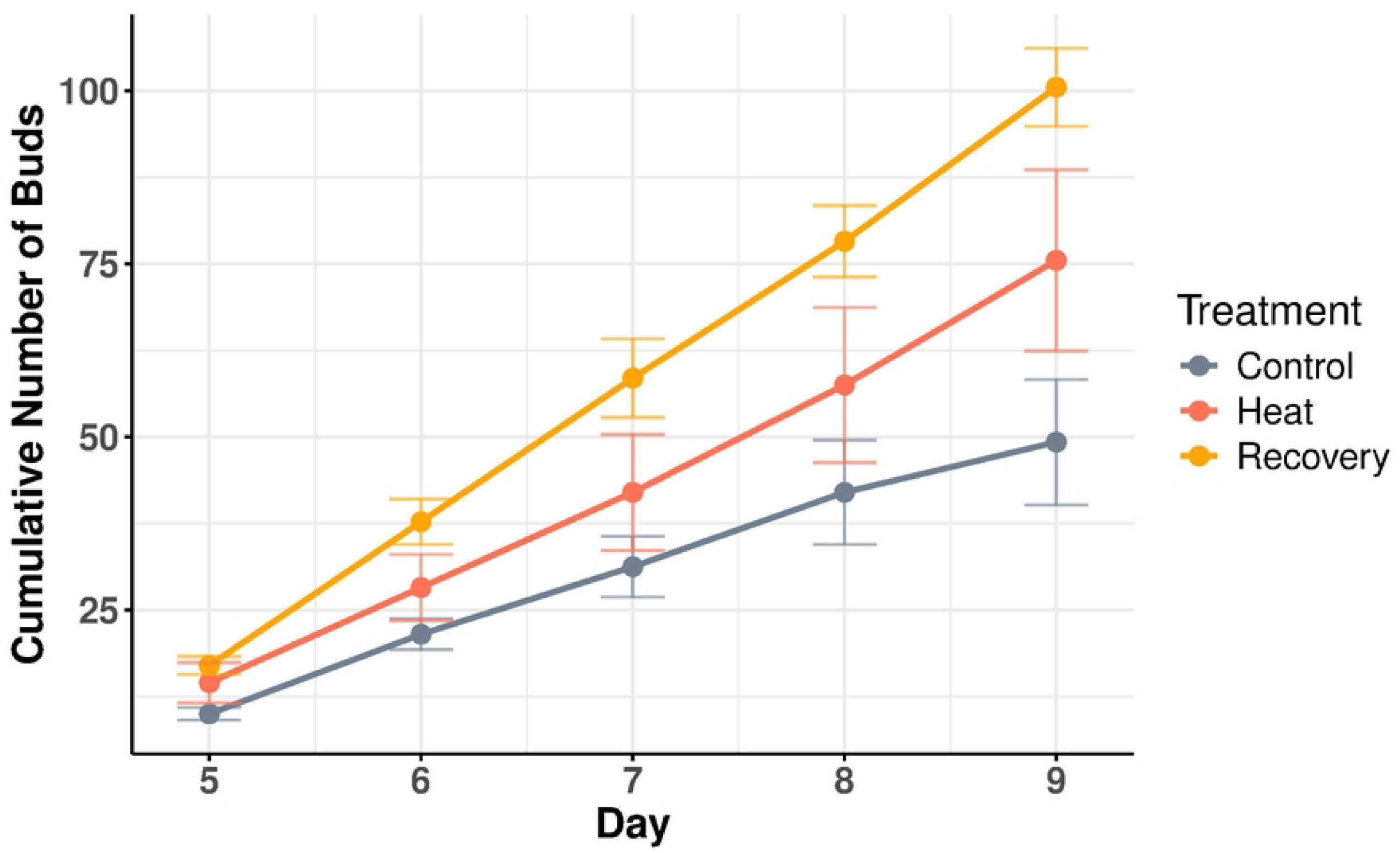
Asexual reproduction via budding over time in each treatment group. Polyps infected with heat-stressed symbionts (heat and recovery groups) produced more planuloid buds compared to the control.

Many jellyfish are capable of rapidly proliferating and blooming in response to rising temperatures (14,32). In addition to increased asexual reproduction, they may also shift from asexual reproduction to strobilation, presumably to escape unfavorable conditions and facilitate habitat expansion through sexual reproduction (32). However, in *C. xamachana*, strobilation is tightly linked to the presence of a specific photosymbiont, potentially limiting its capacity to respond quickly to environmental change unless a compatible algal partner is present. Why, then, would photosymbiotic jellyfish evolve reliance on symbionts for such a key life history transition? This dependency raises the possibility that symbiont establishment functions as a developmental checkpoint, ensuring that environmental conditions can support a stable symbiotic association, and thus medusa survival and dispersal. Supporting this idea, microbes are known to act as early indicators of environmental shifts in animal hosts (33), and animals have evolved systems like the innate immune response to detect and respond to microbial cues. These immune pathways have been co-opted for developmental regulation in some specie (34) and may underlie the link between symbiosis and strobilation in *C. xamachana*. Whether this reliance on a photosymbiont ultimately provides an adaptive advantage by aligning symbiosis success with environmental suitability remains to be tested.

## Conclusions

Our study provides evidence that heat-stressed *Symbiodinium* disrupts *C. xamachana* strobilation, delaying metamorphosis and, under sustained stress, completely preventing it. Developmental abnormalities, such as inverted bell morphologies and increased symbiont density, suggest altered host-symbiont dynamics, while increased asexual reproduction indicates a stress-induced survival strategy. These findings highlight the broader implications of thermal stress on cnidarian-algal symbiosis. Future research should investigate the biochemical cues driving strobilation, particularly the role of heat-induced metabolite shifts, molecular mechanisms under the physiological changes, and explore whether similar disruptions occur in other photosymbiotic cnidarians. As climate change continues to drive ocean warming, the breakdown of cnidarian-algal symbiosis may have cascading effects on marine ecosystems. By improving our understanding of how *Cassiopea* and its symbionts respond to heat stress, this study contributes to a growing body of research on the resilience and vulnerabilities of photosymbiotic organisms, specifically on the role of host-microbe interactions driving marine invertebrate development in an era of global climate change.

## Acknowledgments

This work is funded by a Packard Fellowship for Science and Engineering (2019–69653) to JL. We thank Drs. Pieter Johnson and Nicole Lovenduski from the University of Colorado Boulder for their constructive comments on earlier versions of the manuscript.

## Supporting information

**Table S1: Algal cultures chlorophyll t-test data**

**Table S2: Strobilation rate data**

**Table S3: Algal cultures cell count data**

**Table S4: Algal cultures chlorophyll concentration data**

**Table S5: Ephyra algal density data**

**Table S6: Asexual budding data**

